# Predicting single-stranded DNA oligonucleotides 3D structures: an open issue

**DOI:** 10.1101/2025.10.28.684999

**Authors:** Selma Bengaouer, Thomas Binet, Stéphane Octave, Séverine Padiolleau, Bérangère Avalle, Irene Maffucci

## Abstract

Single-stranded Nucleic Acids (ssNAs) play major biological functions and represent an interesting biotechnological tool. Their function depends strictly on the specific foldings they can adopt. Therefore, information about ssNAs’ tridimensional structures is fundamental to investigate their functions. In this context, *in silico* 3D structure prediction can facilitate ssNAs design. Many algorithms have been developed with this aim, mainly focused on RNA. However, the growing interest in single-stranded DNA (ssDNA), due to their greater stability as compared to RNA, has highlighted the need to adapt these methods for ssDNA. This study assessed three RNA 3D structure prediction methods (RNAComposer, SimRNA, and Vfold3D), based on their performance in the Critical Assessment of protein Structure Prediction 15 and/or 16, to evaluate their applicability to ssDNA. At this scope, a dataset of 93 experimentally determined ssDNA structures, including challenging motifs such as G-quadruplexes, was built. Various metrics, such as RMSD, GDT TS, and INF were employed to benchmark the accuracy of the predictions. The three tools showed similar and moderate performances. In addition, they show strong difficulties in modeling G-quadruplexes, and structures containing motifs strongly increasing the intrinsic flexibility of ssDNA. Despite the recent efforts in the prediction of the 3D folding of ssNAs, it is clear that a significant improvement of the methods is needed. This should involve taking into account the conformational variability of this kind of molecules and paying attention to their specific 3D motifs.

**Author summary:** Single-stranded oligonucleotides (ssNAs) are RNA or single-stranded DNA molecules involved in crucial biological processes, such as gene expression, DNA replication, and transcription. In addition, they represent a powerful biotechnlogical tool exploitable in therapeutics, diagnostics and biosensing. This is due to their capacity of recognizing different kind of molecular targets, thanks to the tridimensional foldings they can adopt. It is therefore clear that the knowledge of the ssNAs structure is fundamental to master these molecules. With this aim, much effort has been paid in developing computational tools for the prediction of ssNAs 3D structures. However, their application is mostly limited to RNA sequences, even if the interest in ssDNAs is rapidly increasing. Moreover, an extensive benchmark on their performances is missing. We focused this work on assessing the performances of three ssNA structure prediction tools, which best performed in the two latest Critical Assessment of protein Structure Prediction contests, on a large dataset of ssDNA, in order to establish the strengths and limits of the available tools. This knowledge will be helpful in finding new solutions for better understanding the ssNAs folding and their structure-function relationship.

## Introduction

Single-stranded nucleic acids (ssNAs), i.e. single-stranded DNA (ssDNA) and RNA oligonucleotides, play an essential role in biological processes, making them a powerful biotechnological tool [1–3]. ssNAs are highly flexible and can fold into specific 3D conformations through intramolecular base pairing, which are crucial for their stability and function [4]. Because of their intrinsic flexibility, ssNAs structures are characterized by a large variety of base pair networks, named secondary structure motifs, such as hairpin stem-loops, bulges, internal loops, G-quadruplexes, pseudoknots, and multiple way junctions (Fig. 1). These form the basis for higher-order 3D architectures, where additional interactions, including long-distance contacts and molecular stacking, further refine the stability and functionality of ssNAs [5]. This structural richness enables them to bind a wide range of target molecules with remarkable specificity and affinity, achieving dissociation constants in the nano-to picomolar range [1, 2, 6].

**Fig 1.**
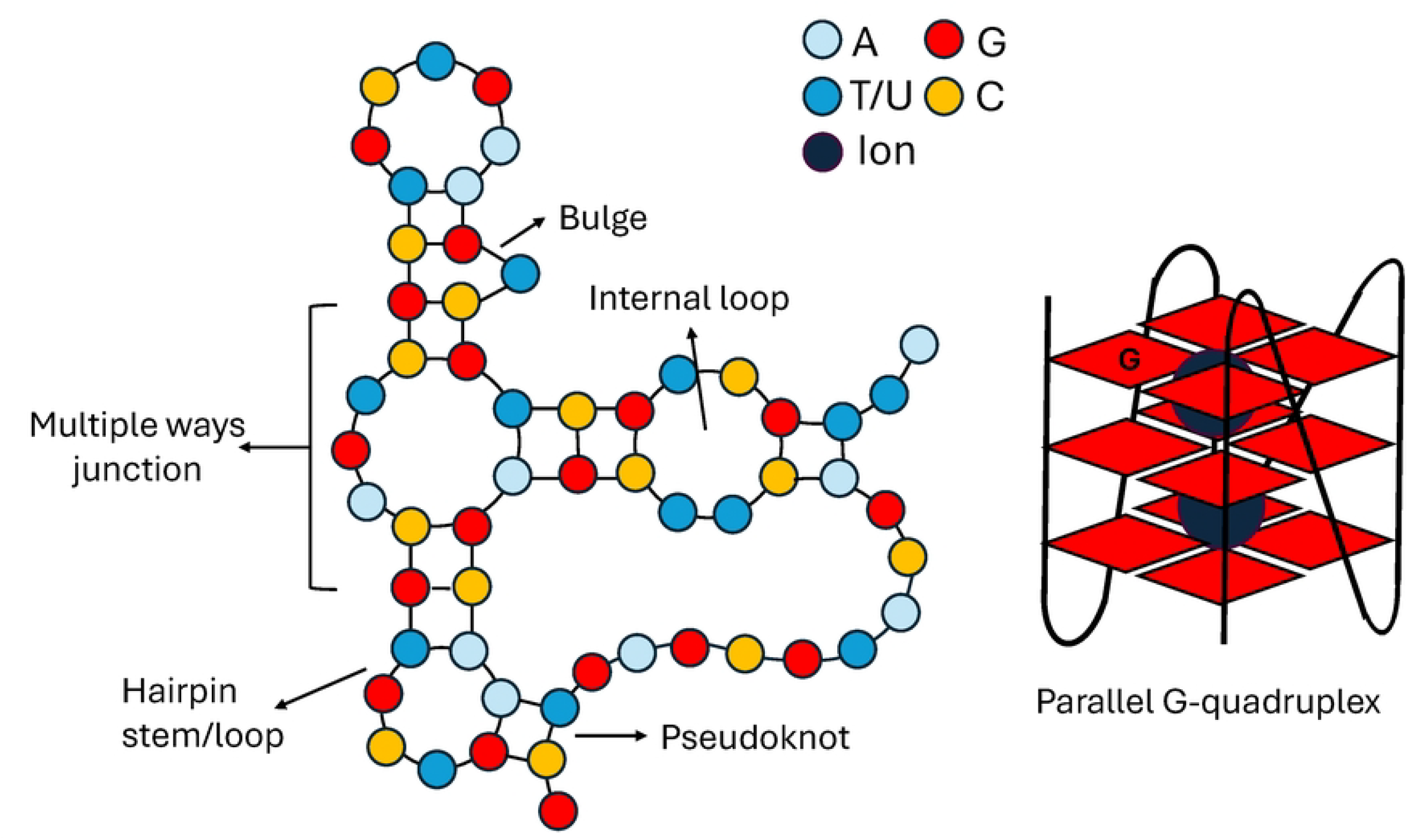
**Main motifs found in single stranded DNA.**

Therefore, it is clear that a high-resolution structural determination is crucial for understanding the biophysical properties and functional mechanisms of ssNAs. However, the characterization of their 3D structures at an atomic resolution remains a significant challenge. Indeed, experimental techniques exploitable at this scope are often complex, expensive, and time-consuming [3, 6–8]. Therefore, computational methods for ssNAs 3D structure prediction provide a promising alternative, generating valuable insights that complement experimental findings. As a consequence, similarly to the protein folding problem [9], the 3D structure prediction of ssNAs is becoming a bioinformatics hot topic, to the point that this kind of molecules have been included in the latest Critical Assessment of Structure Prediction (CASP) challenges [10, 11].

Indeed, the 15th and the 16th editions of the CASP contest [10, 11] included the evaluation of the performances of more than 40 methods for the prediction of RNA 3D structures. However, only a limited number of structures were considered within the CASP15 and CASP16 and the overall results show that the performances of these algorithms are still very far from those of the methods for the 3D structure prediction of proteins.

In addition, it has to be noted that the prediction capabilities of the considered tools in modeling ssDNA were poorly assessed, despite the growing interest toward this ssNA [12], due to its well known synthesis routine [13] and greater stability as compared to RNA, resulting from the absence of the 2’-hydroxyl group found in RNA, which makes DNA less prone to hydrolysis compared to RNA [14]. Despite of this, no ssDNA was present in the CASP15 blind challenge and only two ssDNA was included in the CASP16, with one being in complex with a protein [11].

To thoroughly assess the modeling abilities of the best CASP15/16 oligonucleotides 3D structure prediction tools, we selected three methods, namely, RNAComposer [15], SimRNA [16], and Vfold3D [17], that rank among the top 5 in the CASP15 and/or CASP16 challenges and are freely available as servers or standalone programs. We evaluated their ability in reproducing the experimental structure of 93 ssDNA retrieved from the Protein Data Bank (PDB) [18] and the Nucleic Acid Database (NDB) [19]. The built dataset included a variety of motifs and ssDNA both in the free state and in complex with proteins, allowing for a comprehensive assessment of structural predictions under different biological contexts.

The predictive accuracy of the aforementioned methods was evaluated using multiple structural assessment metrics, including the heavy atoms Root Mean Square Deviation (RMSD), the Global Distance Test Total Score (GDT TS) [20] to assess overall structural similarity, and the Interaction Network Fidelity (INF) [21] to evaluate the accuracy of predicted molecular interactions. This enabled a comprehensive evaluation of how accurately RNA structure prediction tools can model ssDNA structures, highlighting both their strengths and limitations. Vfold3D was the most proficient at capturing global folds and interaction networks, though it could not generate a model for 15 out of the 93 ssDNAs. By contrast, SimRNA and RNAComposer provided a model for the entire dataset, with SimRNA performing slightly better than RNAComposer. However, all methods encountered difficulties with G-quadruplexes (G4), and with structures containing multiple loops, which strongly increase the intrinsic flexibility of ssDNA. Therefore, these results indicate that much effort has still to be done for the modeling of ssDNA and that their structures need to be investigated considering their conformational freedom.

## Materials and methods

### Single-stranded DNA dataset

The dataset used in this study consists of ssDNA sequences having a published experimental 3D structure retrievable from the PDB [18] and the NDB [19] databases. Sequences containing non-natural nucleotides or corresponding to hybrid DNA and RNA sequences were discarded, together with structures with missing nucleotides and those in complex with small molecules. The resulting dataset comprises 93 ssDNA (Table S1), ranging from 7 to 53 nucleotides and including 26 oligonucleotides possessing a G4 motif and 2 oligonucleotides with a long-distance interaction similar to a pseudoknot. For each ssDNA, the experimental secondary structure was extracted from its 3D structure using x3DNA-DSSR [22]. In the case of ssDNAs containing G4 motifs, a hybrid approach was applied: the G4 secondary structure was first predicted using ElTetrado [23], then integrated with the secondary structure obtained via x3DNA-DSSR to reconstruct the complete and accurate 2D structure.

### 3D structure prediction tools

For the designed benchmark we chose to test the predictive capabilities of ssNA 3D structure prediction tools freely available as standalone programs for Unix/Linux distributions or as free web-servers and that performed the best within the CASP15 and/or CASP16 challenge in terms of RNA structure prediction [10, 11]. The tools satisfying the chosen conditions are Vfold3D [17], SimRNA [16] and RNAComposer [15].

Vfold is a computational method for predicting the secondary and/or 3D structure of RNA molecules. The software consists of two algorithms, Vfold2D and Vfold3D. However, in this study, except when otherwise stated, we focused on Vfold3D, since the experimental secondary structure of each ssDNA was provided as input together with its sequence.

The Vfold3D algorithm uses the motif template assembly method [17] to predict the target 3D structure. The process begins with the identification of structural motifs in the target RNA based on parameters such as sequence length and similarity. These motifs are then searched within the Vfold3D motif template database, which has been built from RNA structures available in the PDB. The database contains a broad range of motif templates, including hairpin stems/loops, bulges, internal loops, multiple way junctions, pseudoknots, hairpin–hairpin kissing loops, and 5’-end 3’-end tail loops. However, if specific motifs are absent or not represented in the database, Vfold3D may not accurately predict the target structure. In such cases, the VfoldLA algorithm [24] is employed. It uses a similar assembly strategy but is based on loop templates and helices instead of motif templates to predict unidentified motifs. The assembled structures may exhibit structural clashes or non-ideal features, such as incorrect bond lengths, bond angles, or torsional angles. To efficiently optimize these structures without excessive computational cost, the algorithm uses the IsRNA model [25]. This model combines a coarse-grained Replica Exchange Molecular Dynamics (REMD) simulations with knowledge-based interaction potentials to refine the all-atom structures generated during the assembly. Therefore, in our study, for each initial 3D structure of our database, REMD simulations were performed using 10 replicas. The simulations were run with default parameters, including a simulation time of 5 ns and 1000 recorded structures per replica. The algorithm generated 5 final predicted structures based on clustering of the low-energy structures of the trajectories.

SimRNA is a stochastic computational method for predicting the 3D structure of RNA molecules. It is built on three main functional components: (i) the representation of the simulated molecule by using a coarse-grained model, (ii) a scoring function used to guide the conformational search, and (iii) a method for the sampling of the conformational space. According to the SimRNA coarse-grain model, every nucleotide is represented by 5 beads: the backbone includes 2 beads, one for the phosphate group and another one for the sugar moiety, while the bases are represented by 3 beads. This representation reduces the number of explicitly represented atoms per residue while retaining the key properties of the RNA chain. The scoring function guiding the conformational search assigns a numerical score to each RNA conformation by calculating various terms that capture the physical and chemical properties of RNA (e.g. the base pairing interactions, the base stacking interactions, the torsional angles, and the distance restraints). Finally, the sampling method employed by SimRNA makes use of the Monte Carlo sampling in either single-thread simulations or replica exchange simulations.

In this study, the parameters recommended by the developers have been used. The bonds, angles, and torsions angles weights that impact the energy of the system are reported in Table S2 of Supplementary Information. Briefly, decreasing the temperature factor from 1.35 to 0.9 facilitates the exploration of different conformations. During the single Monte Carlo simulation, the structure is recorded every 16 000 steps for a total of 16 000 000 iterations, resulting in the acquisition of a total of 1001 conformations, including the initial structure. Only the conformation with the lowest energy was retained for each oligonucleotide for further analyses. Additionally, when mentioned, Replica Exchange Monte Carlo (REMC) simulations were conducted using 10 replicas to evaluate whether this method could explore more conformations than in a single simulation, using the same parameters. After the simulations, we performed a clustering using the output trajectories issued from the different replicas. The clustering tool (version 3.20) does not allow setting a fixed number of clusters; it continues to group structures until it reaches the end of the input data. Therefore, for our analysis, we selected the top five clusters for each oligonucleotide and converted the first frame of each cluster into a PDB file for the further analysis.

RNAComposer is an algorithm implemented on a web-server that can predict the complete 3D structure of RNA molecules. The method relies on assembling short fragments extracted from known structural data, which are then combined to reconstruct the full structure. The algorithm uses the RNA FRABASE [26, 27] database to identify and locate the building blocks corresponding to the previously identified short fragments. To use of this algorithm, the default parameters were employed. Notably, 800 structures are initially generated and minimized through 20 000 Monte Carlo cycles. The resulting structures are then ranked by energy and the top 10 are conserved.

For the 3 selected tools, the experimental secondary structure has been provided as input in order to evaluate the predictions quality starting from a unique and experimentally correct base pair network.

### Optimization of the predicted 3D structures

All the selected methods are designed to predict RNA structures on the basis of a RNA input sequence. Vfold3D and RNAComposer can also process DNA sequences by automatically converting them into RNA structures before proceeding with the prediction step, while SimRNA requires as input a RNA sequence. In all cases, the output is in RNA format. Therefore, the obtained RNA structures were manually converted into ssDNA structures by mutating the uracil bases into thymine bases using the PyMol mutagenesis plug-in and replacing the hydroxyl group in C2’ by a hydrogen atom.

In addition, in some cases, the obtained 3D structures showed ruptures in the sugar-phosphate backbone and/or the RNA-to-DNA conversion caused steric clashes due to the greater steric hindrance of the thymine as compared to the uracil. Therefore, a structural minimization was performed in the gas phase using the Amber20 package. This involved the minimization of the hydrogens through 4 000 steepest descent cycles followed by up to 1 000 conjugate gradient cycles. The whole structure was then successively minimized over 16 000 steepest descent cycles and up to 4 000 conjugate gradient cycles.

### Comparison 3D structure metrics

To assess the efficiency and accuracy of the selected tools, each predicted ssDNA tertiary structure was compared to the corresponding experimental reference structure. Various metrics were used to evaluate both global and local accuracy, in agreement with the guidelines of the CASP15 international challenge [10].

In the light of this, we computed the Root Mean Square Deviation (RMSD) as

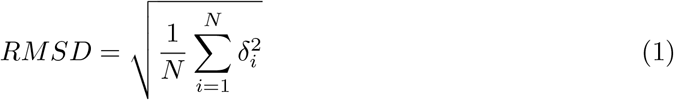

where *δ*_*i*_ corresponds to the distance of the atom *i* of the predicted structure from the same atom in the reference one. In our case we considered the RMSD of the heavy atoms of the predicted structure from the same set of atoms of the experimental structure. In accordance with the literature [28, 29], we considered that a prediction was close to the reference structure when the RMSD was lower than 5 Å.

The Global Distance Test Total Score (GDT TS) was calculated using the LGA tool [20]. This metric evaluates the percentage of C4’ atoms in the predicted structure that fall within predefined distance thresholds of 1 Å, 2 Å, 4 Å, and 8 Å. These thresholds indicate the level of structural similarity between the predicted and reference structures. The GDT TS is derived by summing the percentages calculated for each threshold and it ranges from 0% (no similarity) to 100% (perfect match) (Equation 2). We considered that the prediction globally reproduces the experimental folding when the GDT TS is greater than 45% [30].

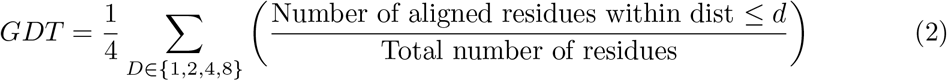

The total Interaction Network Fidelity (INF) score [21] was incorporated in this study to evaluate how well a predicted molecular structure preserves the network of interactions present in a reference structure. The total INF score is computed as:

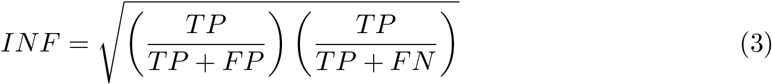

where *TP* (true positive) is the number of interactions (the canonical and non-canonical base pair and the base-stacking, in this study) correctly predicted in the model that correspond to those in the reference structure. *FP* (false positives) is the number of interactions predicted in the model that do not have a corresponding match in the reference structure. *FN* (false negative) is the number of interactions present in the reference structure but missing from the predicted model. The INF score ranges from 0 (low fidelity, the reconstructed network does not accurately reflect known interactions) to 1 (high fidelity, the reconstructed network captures a large proportion of the interactions present in the reference network), with values ≥ 0.7 indicating a globally satisfying prediction of the interaction network [31].

### Secondary structure comparison metrics

The secondary structure of each predicted model was compared to the experimental one. This required, as a first step, the conversion of each predicted 3D structure into its corresponding 2D representation in the dotbracket notation using the standalone version of x3dna-dssr [22]. As previously mentioned, for the structures containing G4 motifs, a combined strategy using both x3dna-dssr [22] and Eltetrado [23] was used. The resulting 2D structure were then compared to the experimental 2D structures initially provided to the algorithms.

The comparison was made using AptaMat [32] as metric, a highly sensitive matrix-based algorithm designed specifically for comparing aligned secondary structures of single-stranded nucleic acids (ssNAs). An AptaMat score of 0 indicates that the two compared secondary structures are identical, an AptaMat score ≤1.5 refers to close secondary structures, and an AptaMat score *>* 1.5 suggests that the two compared secondary structures differ [33].

## Results and Discussion

To investigate the strengths and weaknesses of the available ssNA 3D structure prediction tools and their applicability to ssDNA, we selected three existing methods developed at this scope that offer a freely available version (detailed in Materials and Methods section) and that were among the best performers in the CASP15 [10] and/or CASP16 [11] challenges: RNAComposer [15], SimRNA [16] and Vfold3D [17]. We tested their prediction performances on a dataset of 93 experimentally determined ssDNA structures with sequence lengths ranging from 7 to 53 nucleotides (Table S1, Supplementary Information). The dataset comprises all kinds of structural motifs (Fig. 1), including G4 and pseudoknots, in order to extensively challenge the chosen tools. To further challenge the selected prediction tools, ssDNA whose structure was resolved in a free form or in complex with a protein was included in the dataset. Indeed, ssNAs are characterized by a set of conformations in dynamical equilibrium, which can be shifted and perturbed by interaction with another molecule [34]. It is therefore interesting to investigate how the ssNA structure prediction tools behave in this scenario.

The three selected prediction tools are developed to model RNA structures. Therefore, a RNA to DNA conversion is required before assessing the quality of the modeled structures (see the Materials and Methods section for details). Successively, each model was compared to the experimental structure by using different metrics commonly used to collate oligonucleotide 3D structures, namely the heavy atoms RMSD, the total INF score and the GDT TS, which can provide additional information about the quality of their topologies (Table S3). To facilitate the analysis and discussion of the results, for each metric, a quality threshold was fixed according to the literature [21, 28–30]. More precisely, a good quality model should have a heavy atoms RMSD ≤ 5 Å from the experimental structure, an INF score ≥ 0.7, and a GDT TS ≥ 45%.

### Global analysis of the prediction tools performances

First of all, it has to be noted that RNAComposer and SimRNA were able to successfully generate 3D models for the whole ssDNA dataset, while Vfold3D failed to produce 15 models out of 93 structures (Table S4, Supplementary Information). These 15 structures include 7 ssDNAs containing a G4 motif and 8 very short sequences (ranging between 7 and 14 nucleotides). They share a common characteristic: their secondary structure is characterized by a single isolated base pair, making the folding prediction particularly challenging. As a consequence, the overall performance of the Vfold3D tool is slightly misestimated due to its failure to predict 24% of the dataset.

Besides this, the three prediction tools provide medium quality and overall equivalent results, with none of them excelling in predicting the ssDNAs 3D structures and, at the same time, completely failing in the task (Fig. 2). More precisely, all the methods tested in this study show relatively similar median heavy atoms RMSD values, 7.4 Å, 6.7 Å, and 6.6 Å for RNAComposer, SimRNA, and Vfold3D, respectively, which are all above the threshold of 5 Å (Fig. 2a). Notably, Vfold3D achieves the lowest median RMSD, demonstrating the relatively high accuracy of this tool. Furthermore, 36% - calculated over the 78 structures Vfold3D handled - of the structures predicted by Vfold3D have a RMSD below 5 Å. This suggests that Vfold3D is effective in generating accurate structural models, when it is able to provide a prediction for a given sequence and secondary structure. Moreover, this tool is the one generating the models with the lowest RMSD values, as shown in Fig. 2a and Fig. S1. Nevertheless, these results need to be rightsized because of the previously mentioned limits of this tool in finding a solution for particular structures. In addition, although showing higher median heavy atoms RMSD, both RNAComposer and SimRNA provided a similar amount, 32% and 31%, respectively, of models with a RMSD lower than the threshold, with these percentages being on the whole dataset. Interestingly, SimRNA is the tool providing the narrowest RMSD distribution for its models, with no model having a RMSD above 17.5 Å (Fig. 2a).

**Fig 2.**
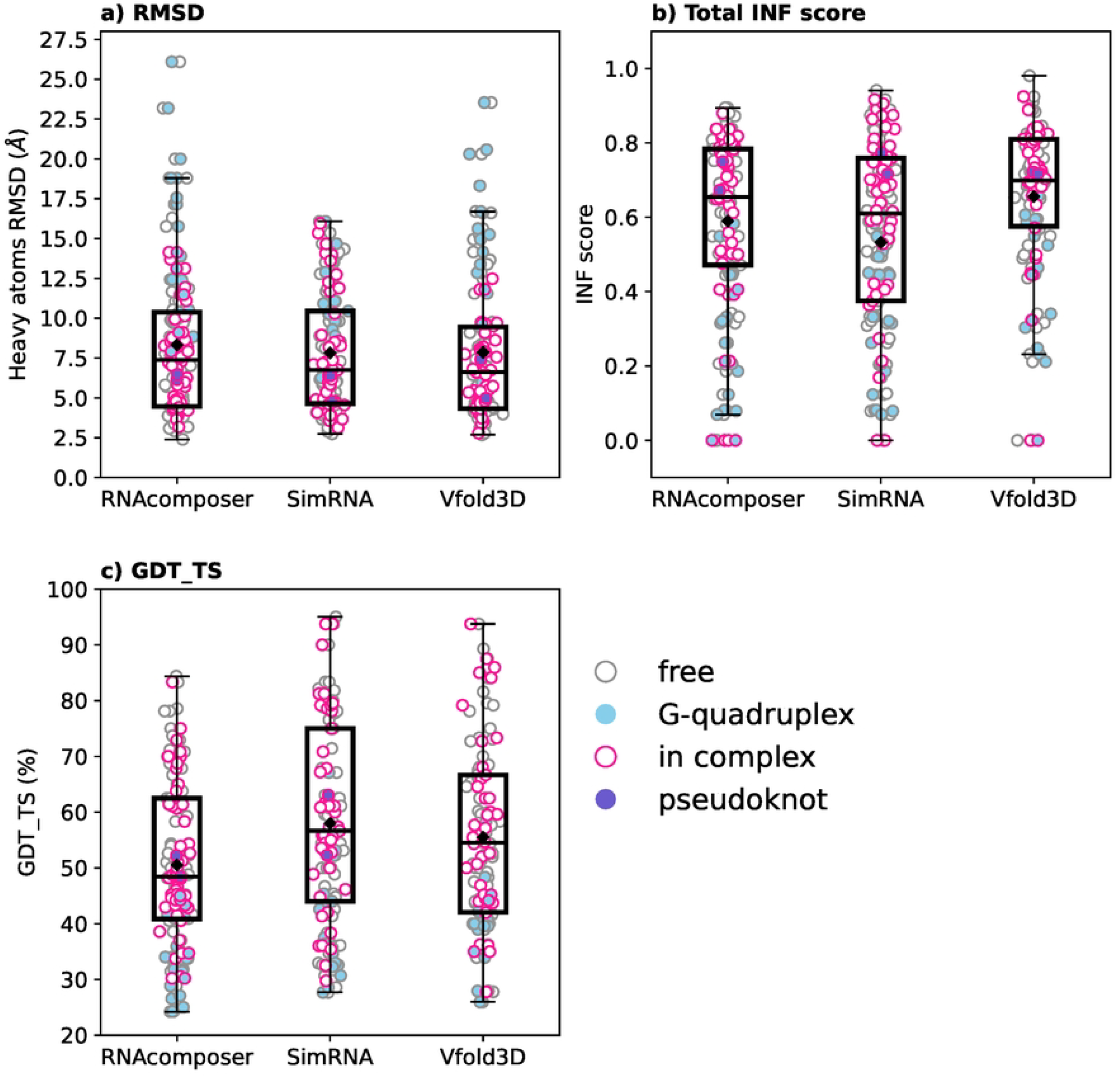
**Boxplots of (a) heavy atoms RMSD, (b) INF score, and (c) GDT TS obtained for the models provided by RNAComposer, SimRNA and Vfold3D for the 93 ssDNA structures. The mean values are indicated as black diamonds; the G-quadruplexes and the pseudoknots are indicated as light blue and purple dots, respectively, while the contour of the dots indicates the ssDNA state: free (gray) and in complex with a protein (magenta).**

However, the RMSD has notable limitations, such as its dependence on the molecule size, its sensitivity to the structures alignment, and its difficulty in capturing local details, highlighting the need for additional complementary metrics. Therefore, we selected two additional metrics: the GDT TS, and the total INF score. This latter allows to investigate the quality of the interaction network (all kind of base pair and base stacking) of the model as compared to the reference structure. Fig. 2b shows the results obtained for this metric by the 3 prediction tools. As seen for the RMSD, Vfold3D achieves the best median INF score (0.71) and the highest amount (53% over 78 structures) of predictions above the 0.7 threshold, suggesting that it is more performing in maintaining the key structural contacts as compared to the other methods. Nevertheless, the fact that 15 of the 93 structures could not be included in this statistical analysis means that the results may be slightly inaccurate. Interestingly, RNAComposer performs slightly better than SimRNA on the base of the total INF score, with a median value of 0.65 (0.61 for SimRNA), a percentage of predictions above the fixed threshold of 41% against 34% of SimRNA, and a narrower score distribution. This indicates that, although RNAComposer may have higher RMSD values, it is able to reproduce the interaction network of ssDNA slightly better than SimRNA (Fig. 2b).

The GDT TS score is used to assess the global similarity between a model and a reference structure. *Vis-à-vis* of this metric, RNAComposer exhibits the lowest performance, with a median GDT TS of 48.4% (Fig. 2c) and 60% of the predictions above the threshold of 45%. This, together with the previous results, indicates that RNAComposer can provide models with interactions quite close to those of the reference structure, but suffers from limits in providing a correct spatial organization. On the other hand, SimRNA and Vfold3D show better and similar results, with median values of 58.1% and 56.9%, respectively, and a percentage of models above the threshold of 76% and 73%, respectively. However, a more detailed analysis of the GDT TS distributions reveals a slight advantage for SimRNA (Fig. S1). However, once again, the Vfold3D statistics being on 78 structures instead of 93 structures, its performance may be misestimated.

The Vfold3D failure to model 15 ssDNAs out of the 93 ssDNAs likely originates from its inability to find compatible motifs that meet the base pair constraints in the experimental secondary structure. This may especially occur when a structure contains only a single isolated base pair interaction. Indeed, if, instead of providing the experimental secondary structure constraints as an input, the whole Vfold procedure is applied (i.e. the use of Vfold2D for the prediction of the secondary structure followed by the modeling of the 3D structure prediction with Vfold3D), Vfold returned a model for 14 out of 15 sequences (Table S5, Supplementary Information). Nevertheless, these models are based on a Vfold2D-predicted secondary structure which not fully corresponds to the experimental one. As a consequence, all the modeled structures have INF scores ≤ 0.7, implying that the models fail to accurately reproduce the native interaction networks (Table S6, Supplementary Information). If we consider the other two metrics, the very short ssDNA (≤ 10 nucleotides) without a G4 motif (2A0I, 5GWL, 5GWQ, 5OND, 6J37, 6M0B, and 6M0C) display a good structural similarity with native forms, as indicated by GDT-TS scores ≥ 45% and RMSD values ≤ 5Å or close to the threshold. For example, in the case of the minidumbbell (PDB code 6M0B [35]), Vfold3D fail to provide a 3D model when the experimental base pair constraints were given as input, but it was able to predict a 3D structure when no constraints were added and we let Vfold2D suggest a secondary structure (.((..)).), which differs from the experimental one ((..)(..)). The experimental structure contains two isolated base pair, which is difficult to detect and model. As a consequence, the experimental interaction network was not reproduced in the 3D model resulting from the whole Vfold pipeline, since no correct interactions were detected (INF = 0.0). However, because of the small size of this ssDNA, the predicted structure showed a heavy atoms RMSD of 5.0 Å and a GDT TS score of 68.7%.

All the other structures (2KF8, 2N21, 5F55, 5NYS, 5VHE, 6SUU, and 6T2G) are all but one characterized by the presence of a G4 motif, and, together with a low INF score, their models show a poor GDT-TS (≤ 45%) and a high RMSD (≥ 5Å), suggesting inaccurate global folding.

The only ssDNA that remained unmodeled even when using the whole Vfold pipeline is a 24 nucleotide oligonucleotide composed of eight GGA triplet repeats (PDB code 1OZ8) [36]. This particular type of repeat is known to adopt a wide variety of structures [37–41]. In the case of 1OZ8, it forms a distinct structural motif characterized by an intramolecular higher order tail-to tail packing of parallel G4. This particular folding pattern likely contributes to the Vfold inability to generate either a 3D model or even a secondary structure for this ssDNA.

The failure in modeling structures with isolated base pair appears to be specific to Vfold, as it was not observed with the other algorithms. Both SimRNA and RNAComposer were able to generate models for these structures, though in the case of 1OZ8 the obtained models were very far from the peculiar experimental 3D structure, with RMSD values of 14.7 Å and 18.8 Å, INF scores of 0.12 for both, and GDT scores of 30.7% and 25.0%, respectively.

### The effect of oligonucleotides length

In order to provide a more detailed information about the performances of the three prediction tools, the dependence of their prediction accuracy from the ssDNA length was assessed. With this aim, the Spearman correlation between the sequence length and the three structural metrics (RMSD, INF, and GDT TS) was computed. The results reported in Fig. 3 and in Fig. S2 show that the INF score does not correlate with the sequence length for any of the three tools, indicating that the tools ability in reproducing the interaction network does not depend on the ssDNA size.

**Fig 3.**
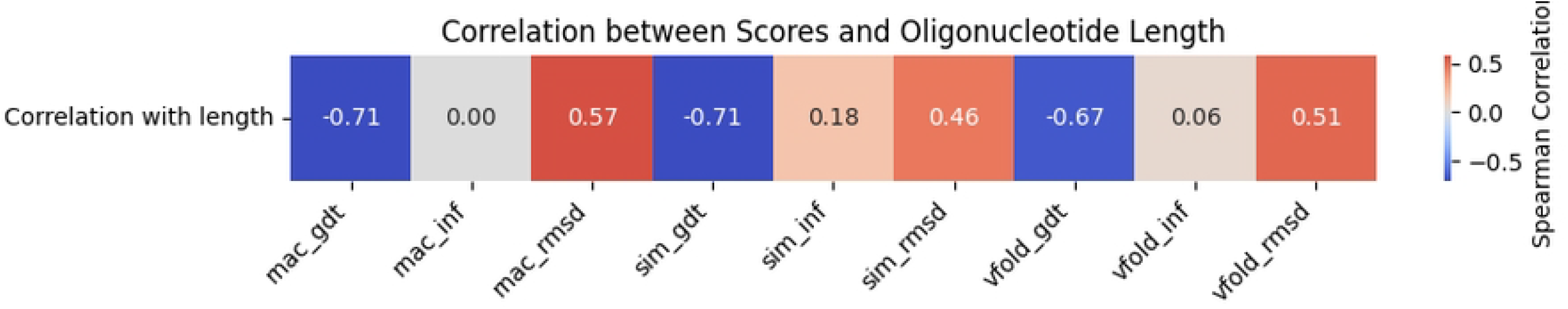
**Correlation between the three structural metrics (RMSD, INF and GDT TS) across the three prediction tools (RNAComposer, SimRNA and Vfold3D) and the oligonucleotide length**

Conversely, the RMSD and GDT TS are moderately correlated with the sequence length across all tools (Fig. S2 and Fig. S3). The heavy atoms RMSD exhibits a weak positive correlation with the ssDNA length (RNAComposer: *ρ* = 0.56; SimRNA: *ρ* = 0.46; Vfold3D: *ρ* = 0.45). A positive correlation would indicate that the longer the sequence is, the higher the RMSD of the provided model is, thus the lower the structural accuracy is. However, the obtained poor correlation mostly depends on the heavy atoms RMSD obtained for very short oligonucleotides (*<* 20 nucleotides). Indeed, as previously mentioned, a low RMSD is expected for short sequences. If these short ssDNAs are removed, the weak correlation disappears, with a *ρ* = 0.18, −0.03, and 0.13, for RNAcomposer, SimRNA and Vfold3D, respectively.

The GDT TS correlation to the ssDNA length is more marked as compared to the RMSD one, with a mild negative correlation, *ρ*, of −0.71 for RNAComposer and SimRNA, and −0.67 for Vfold3D, indicating that the models obtained for long sequences have low GDT TS, which are indicative of a poor quality structure as compared to the experimental one.

In the light of these results, a clear correlation between ssDNA size and modeling abilities of the three tools cannot be determined.

### The issue of the G-quadruplex motif

Giving a closer look to the obtained results, it is possible to highlight some evident limits common to the three tools, with the most noticeable one being the modeling of ssDNAs containing G4 motfs. This issue is brought to the extreme by Vfold3D, which did not provide a model for 7 out of 26 G4-containing ssDNAs. Indeed, if these 26 structures are removed from the dataset, all the metrics improve (Table S7,

Supplementary Information), with medians close to (RMSD) or above (INF and GDT TS) the fixed thresholds.

More in details, for all the methods, none of the models containing a G4 motif had a RMSD ≤ 5 Å or an INF score ≥ 0.7. In addition, only 2 (2M8Z and 5CMX), 11 (2M91, 7CV3, 1I34, 1HAO, 148D, 7CV4, 6EVV, 5CMX, 2M8Z, 2M90 and 5VHE), and 3 (2M91, 6EVV and 2M8Z) structures predicted by RNAComposer, SimRNA, and Vfold3D, respectively, provided a GDT TS ≥ 45%. This slightly better performance suggested by the GDT TS can be attributed to the fact that it uses a more nuanced approach to assess structural similarity by considering a range of distances between corresponding atoms in the predicted and reference structures (as detailed in the Materials and Methods section).

Among the G4-containing ssDNA, 2M8Z is the one for which the three metrics provided the best results. It is composed of 27 nucleotides organized in two distinct regions: a hairpin stem/loop and a G4 formed by two G-quartets (Fig. 4). RNAComposer generated a model with a heavy atoms RMSD of 9.9 Å, a total INF score of 0.51, and a GDT TS of 48.1%; the SimRNA model has a heavy atoms RMSD of 6.2 Å, a total INF score of 0.51, and a GDT TS of 67.1%; finally, Vfold3D globally provided the best model, with a heavy atoms RMSD of 6.0 Å, a total INF score of 0.5, and a GDT TS of 58.3%. Nevertheless, the acceptable performances obtained for this structure can be mainly attributed to the accurate prediction of the hairpin stem/loop region. Indeed, when the G4 motif is excluded from the metrics computation, the INF scores improve, with values of 0.87, 0.90, and 0.90 for RNAComposer, SimRNA, and Vfold3D, respectively. Similarly, the RMSD values for the isolated hairpin loop significantly drop below 5 Å for SimRNA and Vfold3D (4.4 Å, and 4.3 Å). The RNAComposer model exhibits a slightly higher RMSD of 5.2 Å which remains very close to the threshold, the difference being probably due to the flexible loop in the hairpin stem/loop motif.

**Fig 4.**
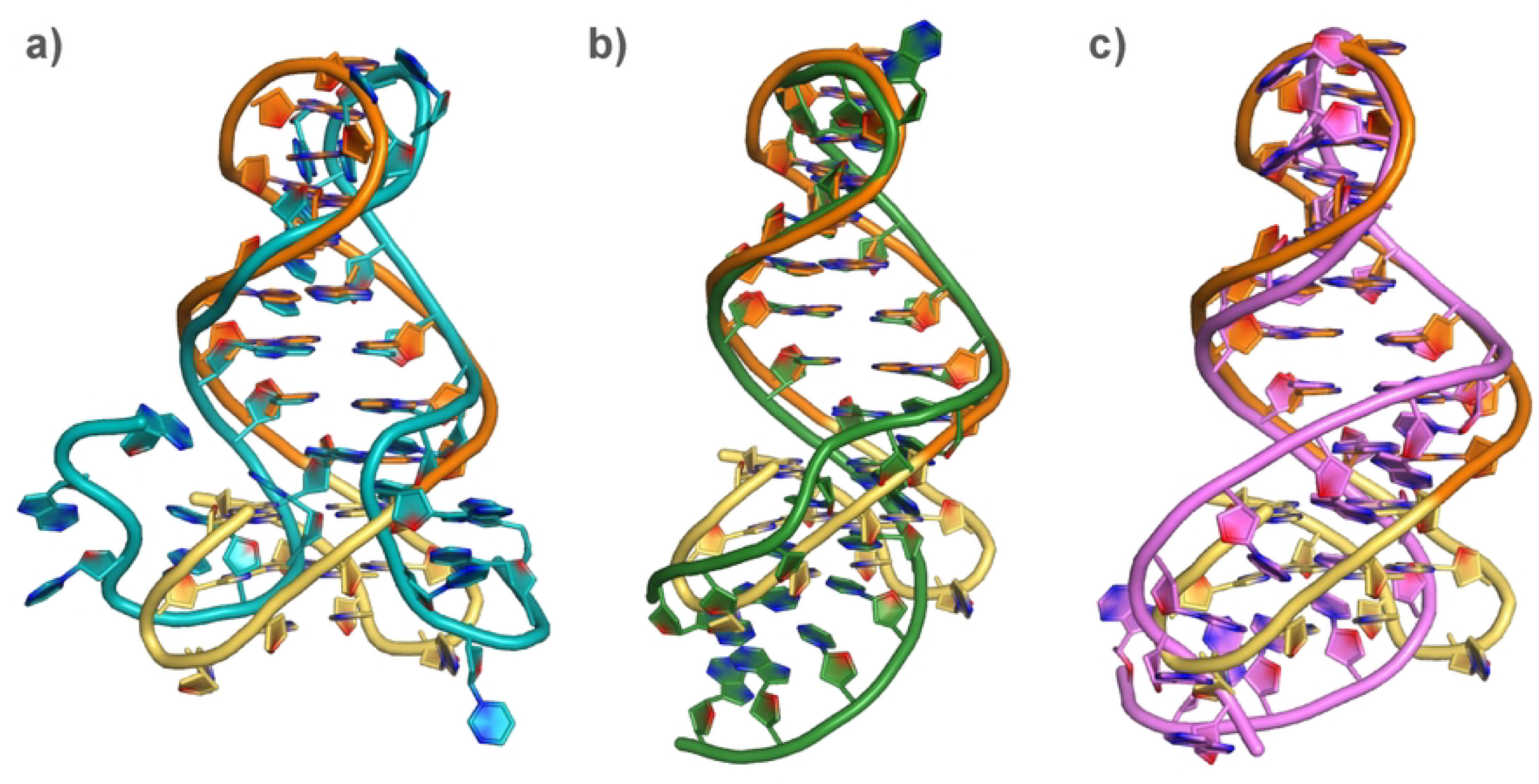
**Alignment of the a) RNAComposer, b) SimRNA, and c) Vfold3D predicted structures to the experimental structure of the 2M8Z ssDNA. The alignment has been realized by focusing on the only hairpin stem-loop motif phosphate backbone. The experimental structure hairpin stem-loop and G-quadruplex are colored in orange and yellow, respectively.**

Even the GDT TS scores, despite being a less stringent metric, improve: 61.8%, 66.28%, and 70.6% for RNAComposer, SimRNA, and Vfold3D, respectively, highlighting a generally higher predictive accuracy in the absence of the G4 motif. Fig. 4 illustrates the quality of the alignment of the hairpin predicted loop when the G4 motif is ignored.

Conversely, the G-quadruplex motif remains more challenging to predict. Indeed, no evidence of the formation of G4 is observed, and not even a partial rearrangement of the guanines into a quartet-like structure typically associated with G4 can be found. These results suggest that the overall predictive accuracy is largely affected by difficulties in modeling the G4 motif. This poor prediction is illustrated in Fig. 4 where we can observe that the experimental G4 (in yellow) is not reproduced by any of the provided models.

The obtained results for the G4-containing structures are not surprising: this motif is known for its intricate topology and stability, with hydrogen bonding on both the Hoogsteen and the Watson-Crick faces, and the presence of monovalent cations to stabilize the structure. The inability to explicitly include ions in the prediction process, together with the atypical base pair characterizing this motif may explain the poor performances of the selected prediction toolsand the failure to accurately capture these interactions. Potentially, the inclusion in the prediction algorithms of a learning step including, among all the data, those about G4 motifs would be an advantage when dealing with these structures. This can be proved by considering the potential performances of AlphaFold3 (AF3) [42] predicting this kind of motif. Although we could not include AF3 in this benchmark because the dataset we used is likely to be part of the AF3 training set and, therefore, all the ssDNAs will be well modeled, we tested it for the structure prediction of a series of aptamers we recently selected against the *Borrelia burgdorferi* CspZ protein [43]. Some of these DNA aptamers were experimentally shown to fold into a parallel G4. This folding is correctly predicted by AF3 by adding an adequate number of K^+^ ions (Fig. S5).

### Dealing with oligonucleotides flexibility

In addition to failing in predicting the G4 motif, the three selected prediction tools significantly struggle when the flexibility of ssDNA is particularly significative, as it is the case, for example, in structures containing loops composed by more than 3 nucleotides (e.g. 1JVE), multiple-way junctions (e.g. 1EZN), unpaired extremities (e.g. 1OSB) and ssDNA in complex with a protein, which may affect the orientation of nucleotides composing the bulges or the loops (e.g. 6SEI).

Although the conformational freedom of an ssNA can be taken into account by the different algorithms during the steps preceding the final structure generation (e.g. SimRNA uses Monte Carlo simulations at this scope), the ssNAs structure prediction tools will provide one or, if specified, a limited number of consensus solutions. This might result in predictions that do not fall within the acceptance range for the chosen metrics.

A clear example of this is represented by the structure with the PDB code 1SNJ [44], which corresponds to a three-ways junction ssDNA resolved by solution NMR. The 12 submitted conformers for this structure indicate a potential flexibility located at the major hairpin stem/loop (G18 to T29, in yellow in Fig. 5a). For this structure, all the tested tools provided models that do not satisfy the requirements for any of the metrics: RNAcomposer, Vfold3D and SimRNA produced a structure with a heavy atoms RMSD of 9.6 Å, 8.3 Å, and 7.3 Å, respectively, an INF score of 0.66, 0.67, and 0.66, respectively, and a GDT TS of 34.7%, 36.1%, and 38.2%, respectively (Fig. 5b-d). However, it can be observed that the main issue comes the wrong orientation of the major hairpin stem/loop (in yellow in Fig. 5) as compared to the experimental structure. This causes a distortion of both the hairpin involving the 5’ and 3’ ends and the second hairpin stem/loop. Thus, although the experimental secondary structure is respected, alterations of the interactions within the ssDNA are observed, as indicated by the total INF score. Indeed, if we exclude from the model evaluation the G18-T29 highly flexible region, the RMSD and GDT TS show a mild improvement: for RNAcomposer, SimRNA and Vfold3D the heavy atoms RMSD drops to 6.9 Å, 3.8 Å, and 5.7 Å, respectively, and the GDT TS increases to 51.0%, 52.0%, and 51.0%, respectively. Conversely, the INF score is not particularly affected, since RNAcomposer, Vfold3D ans SimRNA models excluding the major hairpin stem/loop have a total INF score of 0.66, 0.67, and 0.66, respectively.

**Fig 5.**
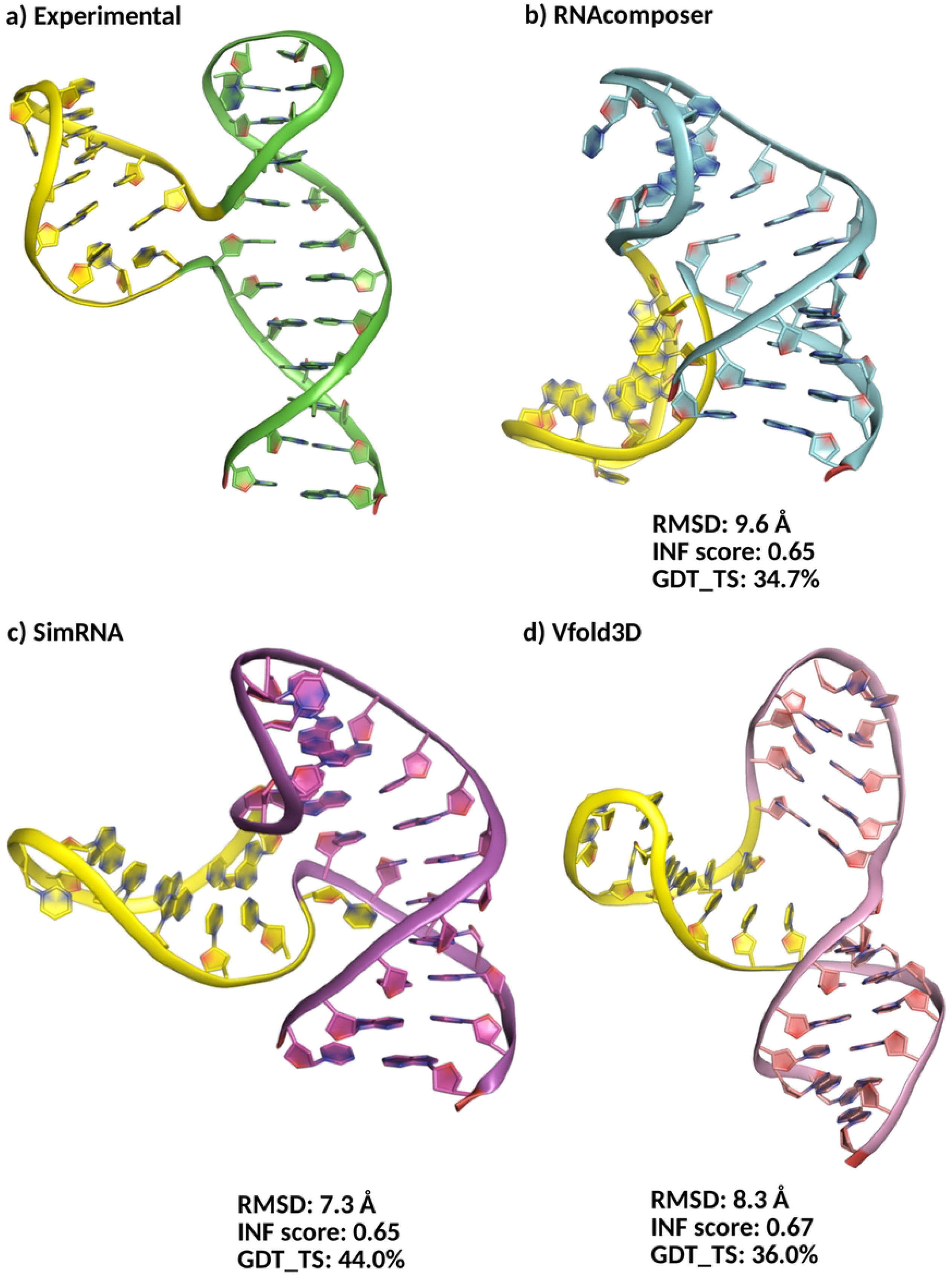
**Experimental (a) and predicted structures (b) RNAcomposer, c) SimRNA, and d) Vfold3D) for the ssDNA with PDB code 1SNJ. The flexible hairpin stem/loop is depicted in yellow. Only the first conformer of the NMR structure is depicted. The heavy atoms RMSD from the first NMR conformer, the INF score and the GDT TS are reported for each model.**

This comparison was initially performed using the first conformer of the 1SNJ structure as the experimental reference. To evaluate whether the choice of conformer could influence the result, we extended the analysis by comparing the predicted structures from each algorithm to all available experimental conformers of 1SNJ (i.e., conformers 2 through 12). However, incorporating these additional conformers resulted in only minor improvements in the structural comparison metrics (Table S8, Supplementary Information), since the flexibility shown by the submitted NMR conformers is limited. For this reason, only the first experimental conformer of 1SNJ was used for the analysis.

Following this, and to better understand how the flexibility of ssDNAs might affect modeling accuracy, we investigated whether the exploration of multiple conformations could improve the results. At this scope, the suboptimal structures generated by RNAComposer, Vfold3D and SimRNA (Table S9, Supplementary Information) were analyzed. SimRNA does not directly produce several models, but, by running a Replica Exchange Monte Carlo (REMC) simulation followed by clustering, five lowest-energy structures from the five most populated clusters could be selected to represent alternative conformations. We pause to note that, since the SimRNA suboptimal structures are produced by using a different simulation approach, the model generated with the default approach will not correspond to the top model obtained with the REMC-based approach. For example, for 1SNJ the SimRNA single model showed a heavy atom RMSD of 7.3 Å from the experimental structure, an INF score of 0.65, and a GDT TS of 44.0% (Fig. 5c). Conversely the top SimRNA-REMC model has a RMSD of 8.8 Å from the reference, a GDT TS of 38.2% and an INF score of 0.66.

The first model generated by each algorithm was not necessarily the most accurate. For RNAComposer, the model 10 outperformed the first, with a heavy atoms RMSD of 9.0 Å compared to 9.7 Å, a total INF score of 0.71 versus 0.66, and a GDT TS of 38.9% instead of 34.7%. Similarly, for SimRNA, the second model yielded improved results, with an RMSD of 7.5 Å (vs 8.8 Å), a GDT TS of 41.0% (vs 38.2%), and an INF score of 0.65 (slightly lower than 0.66) as compared to the first model. In the case of Vfold3D, the fourth model was the closest to the experimental structure, achieving an RMSD of 5.9 Å compared to 8.3 Å for the first model.

The better performances shown by some of the alternative models as compared to the one indicated as the best by the three prediction tools may be attributed to more accurate predictions of the base orientations and improved sugar-phosphate backbone torsion angles, leading to a slightly better overall structural fidelity. Nevertheless, it has to be underlined that the three tools were not able to produce a model close to the experimental structure for this ssDNA.

Another interesting case allowing to investigate the limits of the three selected prediction tools when dealing with ssNAs flexibility is represented by the X-ray resolved ssDNA aptamer of the *Plasmodium falciparum* lactate dehydrogenase (code PDB 3ZH2 [45]). This is a 27 nucleotides distorted hairpin with an internal asymmetrical loop (6 bases on one strand and 1 on the other) and an apical tetraloop, potentially inferring flexibility to the structure (Fig. 6a). Both loops participate to the interaction with the target protein, which, therefore, might stabilize the specific loops orientation observed in the X-ray structure.

**Fig 6.**
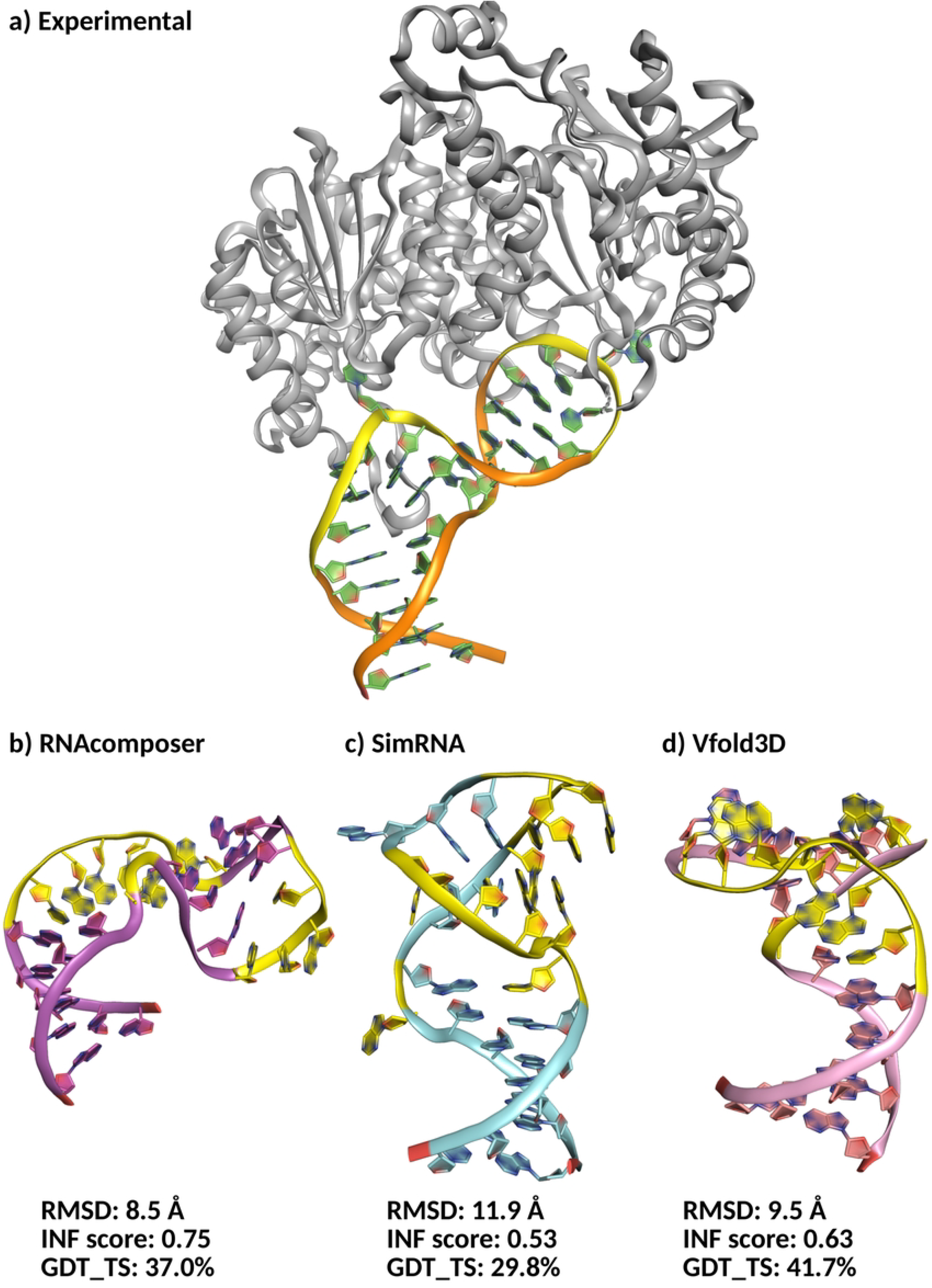
**Experimental (a) and predicted structures by (b) RNAcomposer, c) SimRNA, and d) Vfold3D) for the ssDNA with PDB code 3ZH2. In a) the experimental complex between the ssDNA aptamer and *Plasmodium falciparum* lactate dehydrogenase is reported. The nucleotides which experimentally belong to the flexible loops are highlighted in yellow. The heavy atoms RMSD from the first NMR conformer, the INF score and the GDT TS are reported for each model.**

Similarly to what was observed for the 1SNJ structure, the three 3D structure prediction tools fail in reproducing the experimental conformation, with only the INF score of the model generated by RNAcomposer (0.75) being above the fixed quality threshold (0.7). More precisely, RNAcomposer, SimRNA and Vfold3D provided structures with a heavy atoms RMSD of 8.5 Å, 11.9 Å, and 9.5 Å, respectively, an INF score of 0.75, 0.66, and 0.63, respectively, and a GDT TS of 37.0%, 40.7%, and 44.4%, indicating a global folding far from the experimental one.

Having a closer look at the obtained models (Fig. 6b-d), it is possible to notice that SimRNA and Vfold3D provide a model with a base pair pattern that does not respect the experimental one, indicating that the Monte Carlo sampling implemented by SimRNA and the short REMD simulations performed by Vfold3D fail in dealing with this structure. On the opposite, in accordance with its good INF score, the RNAcomposer model fully respects the experimental base pair, though the orientation of the nucleotides of the two loops impacts also the orientation of the aptamer helices. Conversely to what was observed for the 1SNJ ssDNA, whose structure was resolved as a free oligonucleotide, this difference between the model and the experimental conformations might be due to the fact that the lactate dehydrogenase can stabilize a particular and, potentially, metastable conformation. Indeed, the interaction between the protein and the aptamer can affect the aptamer topology [46], increasing the stability of a conformation not corresponding to the global free energy minimum. If that was the case, the issues encountered by RNAcomposer would be somehow expected and linked to the intrinsic flexibility of the ssDNA and the absence of the protein during the 3D modeling.

As seen for 1SNJ, including suboptimal structures in the comparison did not lead to a significant improvement in prediction results (Table S10). Although some differences are observed between the scores, they remain limited with unmeaningful variations of only *±*1 to 2 Å for the RMSD, 0.1 for the INF, and about 5% for the GDT TS.

It is important to highlight that a ssDNA that has been experimentally resolved in complex with a protein does not represent *per se* an issue for the prediction of its 3D structure. The only limit encountered in this context is connected to the oligonucleotides flexibility and the interaction of the proteins with ssDNA highly flexible regions, such as loops and bulges. Indeed, Fig. 2 shows that the three prediction tools are able to provide good quality predictions *vis-à-vis* of the considered metrics for ssDNAs experimentally available in complex with a protein. For example, for 6 (4KB0, 2VHG, 2A6O, 6FK4, 1UUT, and 4KB1) out of the 46 ssDNA-protein complexes present in the dataset, RNAcomposer, SimRNA and Vfold3D provided models satisfying the threshold values set for the three considered metrics (Table S3). Notably, the ssDNA belonging to these structures are folded into quite simple hairpin stem/loops, with in some cases a bulge of 1 or 2 nucleotides (4KB0, 2VHG, 2A6O, and 4KB1). These ssDNA are, therefore, quite stable, thanks to 5 or more canonical base pairs. More importantly, most of them interact with their target protein through the grooves created by the stem or by their 5’-3’ extremities, while the apical loops are solvent-exposed, meaning that the conformation of the most flexible regions is not impacted by the presence of the protein. 1UUT is the only structure where the apical loop is at the ssDNA-protein interface. However, it is made of only 3 nucleotides, thus its conformational freedom is limited as compared to the previously discussed cases.

### Modeling long distance interactions

Within the built dataset, two structures (5HTO and 5HRU [47]) contain a motif similar to a kissing hairpin pseudoknot, since two hairpins stems/loops form two interactions (Fig. 7a). Both structures correspond to a DNA aptamer, called pL1, folded into a three-ways junction and targeting the *Plasmodium falciparum* lactate dehydrogenase. They differ solely for their crystallization conditions and for a length difference of 2 nucleotides, since the longest pL1 crystallized in 5HTO has an additional T at 5’ and an additional A at 3’. Together with the intrinsic flexibility of the three-ways junction structure, the presence of a long distance interaction, which is also located at the interface with the lactate dehydrogenase, might represent an additional challenge, similar to what previously observed for the flexible ssDNAs in complex with a protein.

**Fig 7.**
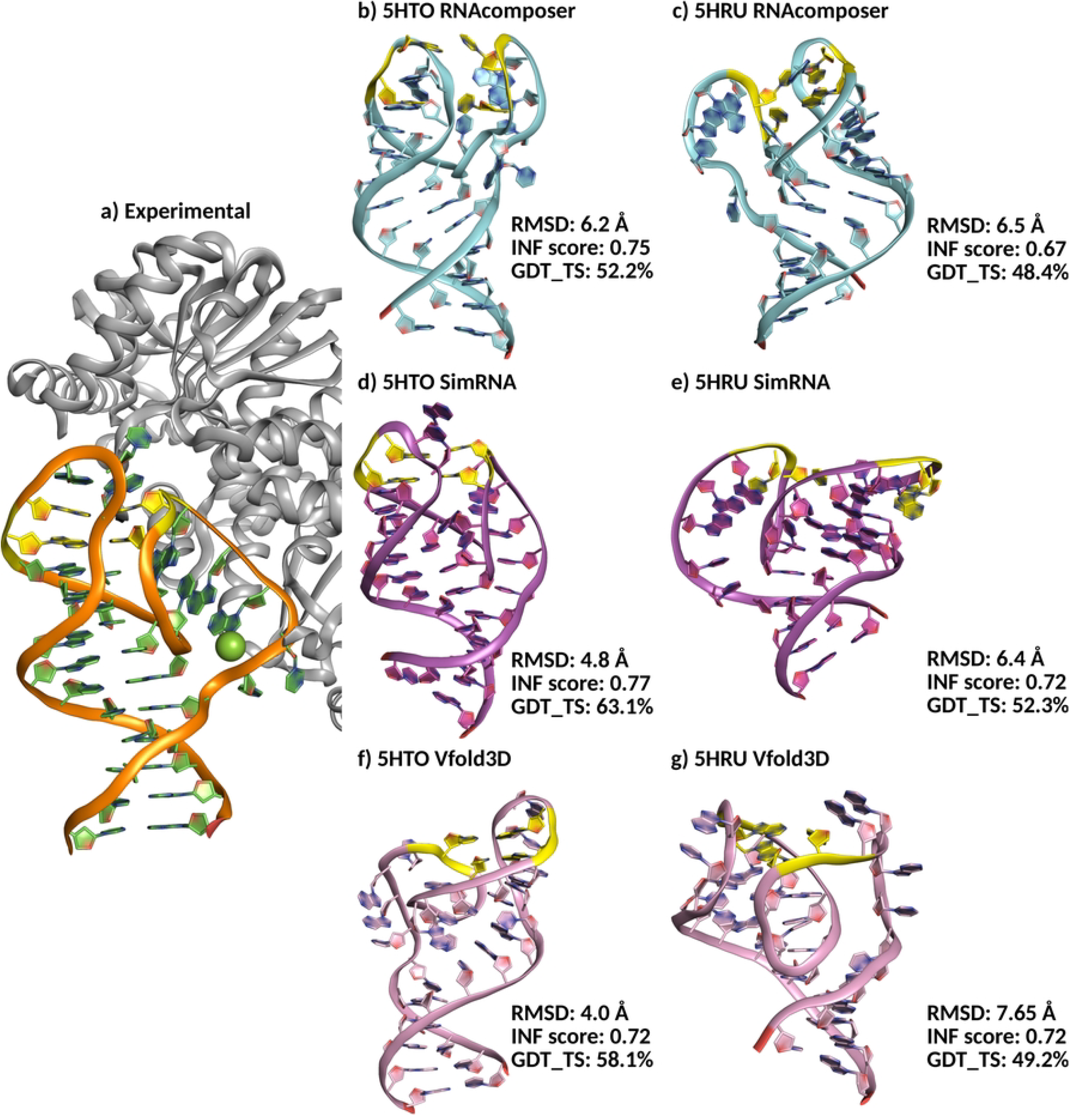
**Experimental structure of pL1 in complex with *Plasmodium falciparum* lactate dehyrogenase (a) and predicted structures for 5HTO pL1 (b, d, and f) and 5HRU pL1 (c, e, and g) by RNAcomposer, SimRNA, and Vfold3D. In a) the experimental complex between the ssDNA aptamer and *Plasmodium falciparum* lactate dehydrogenase is reported. Only the first conformer of the solution NMR structure is showed. The nucleotides which experimentally are involved in the long distance interactions are highlighted in yellow. The heavy atoms RMSD from the first NMR conformer, the INF score and the GDT TS are reported for each model.**

First of all, the three tools successfully provided a model for both structures, where the two kissing loops interact, suggesting that they are able to deal with long distance interactions. Nevertheless, looking at the metrics, a trivial evaluation of the prediction performances concerning this particular case is not possible. Indeed, if we consider the heavy atoms RMSD as compared to the experimental structure, the performances are acceptable only for SimRNA and Vfold3D and only for the pL1 contained in the 5HTO complex, since in the other cases a RMSD *>* 5 Å is obtained (Table S3 and Fig. 7). When focusing on the INF scores, all the tools perform well (INF *>* 0.7) on both structures, except for RNAcomposer that provides a model for 5HRU with an INF score of 0.67. However, this score is close to the fixed threshold value and also to the scores obtained by the other tools (5HRU: 0.72 (SimRNA), 0.72 (Vfold3D); 5HTO: 0.75 (RNAcomposer), 0.77 (SimRNA), 0.72 (Vfold3D)), which means that the three prediction tools are globally equally able to reproduce the nucleotides interactions found experimentally. Finally, by looking at the GDT TS, all the tools provided models with a GDT TS above the fixed threshold, though in all cases the 5HTO aptamer was better modeled as compared to the 5HRU aptamer.

It has to be kept in mind that the three tools got the nucleotides sequences together with the experimental ssDNA secondary structure as input. Despite this, the sole models close to the experimental structure, all metrics considered, are those obtained by SimRNA and Vfold3D for the 5HTO pL1, even if Vfold3D provides a highly distorted orientation of the two hairpins stems/loops forming the long distance interaction. Conversely, RNAcomposer provided a model not respecting the RMSD threshold, but the relative orientation of the kissing hairpins is closer to the experimental one as compared to the one produced by Vfold3D.

For what concerns the 5HRU pL1, the obtained models are of lower quality as compared to the corresponding models provided for 5HTO pL1. Though all the predicted structures show some sort of long distance interaction between the two hairpins stems/loops, the one provided by SimRNA does not involve the two experimentally interacting nucleotides and the orientation of the kissing hairpins of the RNAcomposer and Vfold3D models are not in agreement with the experimental structure. Nevertheless, as previously mentioned, the *P. falciparum* lactate dehydrogenase might have a role in stabilizing the specific conformation observed for this flexible aptamer [47].

Additionally, also in this case, a comparative analysis of the suboptimal structures generated by each method was conducted to assess whether alternative conformations might better reflect the experimental structure. As observed in the previously discussed predictions, the first model is not systematically the most accurate. Moreover, it is interesting to note that the differences between the suboptimal models generated by the same tool are relatively small and do not indicate major structural variation (Table S11 and Table S12).

In the light of these results, it would be interesting to deeply explore the pL1 conformational space, including the Mg^2+^ ion positioned the origin of one of the kissing loops, which seems to be relevant for the correct orientation of this latter. In this context, the selected tools globally would provide good starting point for this task.

### Analysis of the secondary structures of the predicted models

In addition to the 3D structural comparison, an evaluation was conducted to assess how accurately the prediction tools preserved the secondary structure provided as input. For this purpose, the predicted secondary structures were compared to the secondary structures provided as input (see Materials and Methods section and Fig. S6 and Fig. S7). To quantify the differences between the predicted and experimental 2D structures, the AptaMat metric [32] was used. A threshold of 1.5 was fixed based on a previous study [33] to distinguish between accurate and inaccurate predictions, all values ≤ 1.5 are considered close to the experimental secondary structure.

For this comparison, the structures containing G4 motifs were separated from the rest of the dataset, because the previously discussed issue in the prediction of this motif by the three tools implies that the secocondary structure of the generated models will not match the experimental one (AptaMat score *>* 1.5).

Therefore, focusing on the sequences that do not contain a G4 motif or long-distance interactions (67 out of 93 ssDNAs), an overall good conservation of the 2D structure provided as an input was observed. More precisely, RNAComposer achieved a perfect preservation of the experimental base pair pattern (AptaMat = 0) in 73.1% of cases, while Vfold3D and SimRNA showed an AptaMat score of 0 in 44.8% and 41.8% of the cases, respectively. If small deviations from the experimental secondary structure are accepted (i.e. AptaMat ≤ 1.5), RNAComposer provided 98.5%, SimRNA 86.6%, and Vfold3D 79.1% of the models with a secondary structure identical or highly similar to the experimental one. These results can be explained by the algorithm behind each tool. More precisely, RNAComposer builds 3D models directly from a given 2D structure, strictly respecting the base-pair interactions. On the other hand, SimRNA and Vfold3D use more flexible, exploratory approaches, based on REMC and REMD simulations, respectively. Therefore, these latter can potentially modify the base pair pattern given as input. It has to be noted that the satisfaction of the experimental secondary structure constraints does not imply that the final 3D model corresponds to the experimental structure. This underlines the importance of the modeling of loops and bulges, i.e. the fragments without any base pair. These, being highly flexible regions, benefit from the conformational sampling included in SimRNA and Vfold3D, as shown by the better performances of these tools as compared to RNAcomposer.

Concerning the structures containing G4 motifs, as expected, neither RNAComposer nor Vfold3D produced models with an AptaMat score below 1.5. SimRNA generated only one structure with an AptaMat distance below this threshold, corresponding to the structure with the PDB code 2M92. This favorable score can be attributed to the accurate prediction of all the canonical base pairs in the model, which was not the case for the other structures. In addition, where no Watson-Crick base pairs were predicted and only Hoogsteen interactions were observed (e.g. 1HAO), the comparison was not possible. These cases are reported as None values (replaced by −1 in Fig. S7). This failure is mainly due to the absence of a properly formed G4 motif, which prevents its detection and thus the successful 2D reconstruction.

## Conclusion

Single-stranded oligonucleotides are a highly interesting class of molecules, since they are involved in a plethora of biological processes and can be exploited for biotechnological applications. This is due to their ability to specifically and selectively recognize molecular targets, a feature which is strongly related to the three-dimensional structures they can fold into. It is therefore clear that the understanding of the function and usage of these molecules would benefit from the analysis of their 3D structures. At this scope, many *in silico* prediction tools are available and some of them have been evaluated with the CASP15 and CASP16 contests.

In this study, we assessed the performance for the prediction of ssDNAs structures of three 3D structure prediction tools, namely Vfold3D, SimRNA, and RNAComposer, which have been selected based on their performance in the CASP15 and/or CASP16 challenges for the RNA structures prediction. We focused on ssDNA because of their extremely high biotechnological interest in the form of aptamers [1–3] associated to a scarcity of literature on the prediction of their structure.

For this purpose, we built a dataset of 93 experimentally determined ssDNA structures, ranging from 7 to 53 nucleotides, retrieved from the PDB and NDB databases. This dataset includes a variety of structural motifs and ssDNA in both free form and bound to proteins, providing a comprehensive assessment of structural predictions under different biological contexts. The predictive accuracy of these methods has been evaluated using several structural assessment metrics, including heavy atoms RMSD, GDT TS to assess overall structural similarity, and the total INF score to evaluate the accuracy of predicted molecular interactions. The obtained results are in agreement with those reported for the CASP15 and CASP16 contests, which indicate that the ssNA folding problem is far from being solved, since the performances of the structure prediction tools for ssNAs do not reach those shown for the modeling of proteins.

Indeed, the three selected tools have globally similar performances: for RNAcomposer, SimRNA and Vfold3D, respectively, we obtained median heavy atoms RMSD values of 7.4 Å, 6.7 Å and 6.6 Å as compared to the reference experimental structure, median INF scores of 0.65, 0.61 and 0.71, and median GDT TS values of 48.4%, 56.7% and 58.1%. In addition, Vfold3D did not provide a model for 15 out of 93 structures, including 7 G4 and 8 short sequences. This overall suggests that the three tools have medium quality modeling capacities, with satisfactory models obtained for simple structures, such as single hairpins stems/loops or highly stable structures, lacking of particularly flexible motifs (e.g. long loops).

In addition, RNAcomposer, SimRNA and Vfold3D share the same limits. First of all, they performed poorly in modeling G4 motifs: Vfold3D failed to generate 7 out of the 26 G4 structures, and none of the models of the structures containing this motif had an RMSD ≤ 5 Å or an INF score ≥ 0.7. In contrast, three models (6EVV, 2M91, and 2M8Z) satisfy the GDT TS threshold, and among them, model 2M8Z is particularly successful, likely due to its simpler G4 formed only by two G-quartets (instead of the typical three), as well as the presence of more conventional structural elements, such as hairpin stem/loop regions, which facilitate more accurate prediction.

Furthermore, the selected tools struggle to accurately predict the structure of ssDNA molecules that are highly flexible, such as those containing long loops (more than 3 nucleotides), multiple-way junctions, or long unpaired ends. Although SimRNA and Vfold3D try to take into account this flexibility by using methods like Monte Carlo simulations or Replica Exchange Molecular Dynamics (REMD), they still generate only one or a few possible structures. As a result, the predicted models often do not fall within the acceptance range for the chosen metrics. The inclusion of suboptimal models produced by the different tools do not significantly improve the predictions, indicating that the RNAcomposer, SimRNA and Vfold3D exploration of the ssNA conformational space is insufficient.

Finally, the dataset contains two DNA aptamers called pL1 (PDB codes 5HTO and 5HRU) that have a distance interaction similar to a kissing loop hairpin pseudoknot. Importantly, all the tools provided models showing a long distance interaction for the two hairpins stems/loops in contact in the experimental structure. However, only the RNAComposer and SimRNA models for the 5HTO pL1 successfully reproduced the correct relative orientation of the kissing hairpins, while all other models (i.e. including those for 5HRU from all three tools and the Vfold3D model of 5HTO) failed to match the experimental structure. This is potentially due to the presence of a Mg^2+^ ion which might stabilize a particular conformation and/or to the effect of the protein on the pL1 structural organization. Indeed, the hairpin loops interact with the lactate dehydrogenase, which could alter the pL1 conformational dynamics equilibrium in favor of a metastable conformation.

In conclusion, in agreement with what observed for RNA within the two latest CASP constests, RNAcomposer, SimRNA and Vfold3D showed moderate performances in predicting the structure of ssDNAs, with Vfold3D and SimRNA performing slightly better than RNAcomposer. Two main issues remain, the first one being the modeling of G4 motifs-containing structures, and the second one depending on the intrinsic flexibility of ssDNAs. The former limit could benefit from the inclusion of a artificial intelligence-based step during the modeling, which would learn from all the available ssNAs structures, including G4. Conversely, the treatment of the high conformational variability of ssNAs remains more difficult to handle, since any 3D structure prediction tool, by definition, has to provide one or a limited number of final models. Therefore, the exploration of the conformational space of ssNAs still relies on other methods, such as enhanced sampling molecular dynamics techniques, which efficiently allow to do it and to associate a folding free energy and an occurrence probability to each generated conformation.

## Supporting information

**Table S1 Single Stranded DNA dataset**.

**Table S2 Parameters applied to SimRNA algorithm**.

**Table S3 RMSD, GDT TS and total INF scores for the models obtained by RNAComposer, SimRNA and Vfold3D for the ssNAs considered in this study**.

**Table S4 Summary of the structures not predicted by Vfold3D**.

**Table S5 Summary of the structures predicted by the whole Vfold algorithm**.

**Table S6 GDT-TS, RMSD, and INF scores for the 14 predicted structures by the whole Vfold algorithm**.

**Table S7 Statistical comparison of RNAComposer, SimRNA, and Vfold3D on G-Quadruplex vs. No-G-Quadruplex ssDNA structures**.

**Table S8 Comparison of 12 predicted conformers of 1SNJ using RNAComposer, SimRNA, and Vfold3D based on GDT-TS, RMSD, and INF scores**.

**Table S9 Comparison of 1SNJ suboptimal structures predicted by RNAComposer, Vfold3D, and SimRNA based on RMSD, INF, and GDT scores**.

**Table S10 Comparison of suboptimal 3ZH2 ssDNA structures predicted by RNAComposer, Vfold3D, and SimRNA based on RMSD, INF, and GDT-TS scores**.

**Table S11 Comparison of suboptimal 5HTO ssDNA structures predicted by RNAComposer, Vfold3D, and SimRNA based on RMSD, INF, and GDT-TS scores**.

**Table S12 Comparison of suboptimal 5HRU ssDNA structures predicted by RNAComposer, Vfold3D, and SimRNA based on RMSD, INF, and GDT-TS scores**.

**Fig. S1 Value distributions of heavy atoms RMSD, INF score, and GDT-TS obtained for the models provided by RNAComposer, SimRNA, and Vfold3D for the 93 ssDNA structures**.

**Fig. S2 Correlation between INF score and oligonucleotide length for 3D structures predicted by RNAComposer, SimRNA, and Vfold3D**.

**Fig. S3 Correlation between RMSD score and oligonucleotide length for 3D structures predicted by RNAComposer, SimRNA, and Vfold3D**.

**Fig. S4 Correlation between GDT-TS score and oligonucleotide length for 3D structures predicted by RNAComposer, SimRNA, and Vfold3D**.

**Fig. S5 AlphaFold3-predicted 3D structure of an aptamer targeting *Borrelia burgdorferi* CspZ protein, showing the formation of a parallel G-quadruplex motif stabilized by K**^+^ **ions**.

**Fig. S6 Comparison of AptaMat scores between predicted and reference 2D structures for oligonucleotides without G4 motifs, obtained using RNAComposer, Vfold3D, and SimRNA**.

**Fig. S7 Comparison of AptaMat scores between predicted and reference 2D structures for oligonucleotides with G4 motifs, obtained using RNAComposer, Vfold3D, and SimRNA**.

## Acknowledgments

Not applicable

## Notes

### Competing Interest Statement

The authors have declared no competing interest.

